# A benchmarking study of copy number variation inference methods using single-cell RNA-sequencing data

**DOI:** 10.1101/2024.09.09.612120

**Authors:** Xin Chen, Li Tai Fang, Zhong Chen, Wanqiu Chen, Bin Zhu, Hongjin Wu, Malcolm Moos, Andrew Farmer, Feng Zeng, Lijuan Song, Xiaowen Zhang, Wei Xiong, Shusheng Gong, Wendell Jones, Christopher E Mason, Shixiu Wu, Chunlin Xiao, Charles Wang

## Abstract

Single-cell RNA-sequencing (scRNA-seq) has emerged as a powerful tool for cancer research, enabling in-depth characterization of tumor heterogeneity at the single-cell level. Recently, scRNA-seq copy number variation (scCNV) inference methods have been developed, expanding the application of scRNA-seq to study genetic heterogeneity in cancer using transcriptomic data. However, the fidelity of these methods has not been investigated systematically. In this study, we benchmarked five commonly used scCNV inference methods, HoneyBADGER, CopyKAT, CaSpER, inferCNV, and sciCNV. We evaluated their performance across four different scRNA-seq platforms derived data from a multicenter study. We further evaluated the scCNV performance using scRNA-seq datasets derived from mixed samples consisting of five human lung adenocarcinoma cell lines and generated a clinical scRNA-seq dataset from a human small cell lung cancer patient to validate our findings. Our evaluation criteria included sensitivity and specificity of CNV detection, and subclone identification from mixed cancer samples. We found that the sensitivity and specificity of the five scCNV inference methods varied, depending on the selection of reference data, sequencing depths, and read lengths. Overall, CopyKAT and CaSpER exhibited superior performance to other methods, while inferCNV, sciCNV, and CopyKAT outperformed other methods in subclone identification accuracy. Remarkably, inferCNV achieved high accuracy in subclone identification when using data from a “single scRNA-seq protocol”, however, when applying these methods to a dataset derived from multiple scRNA-seq platforms from the mixed samples, we found that batch effects significantly affected the performance of subclone identification for most methods, except for HoneyBADGER. Our benchmarking study revealed the strengths and weaknesses of each of the five scCNV inference methods and provided guidance for selecting the optimal CNV inference method using scRNA-seq data.

## Introduction

Genetic inter-tumor and intra-tumor heterogeneity has been reported extensively in various cancer types^1–6^. Copy number variation (CNV) plays a crucial role in cancer development and progression by amplifying oncogenes or inactivating tumor suppressor genes^7–10^. CNV is an important parameter for characterizing inter-tumor and intra-tumor heterogeneity. While recent studies have highlighted the impact of somatic CNVs on gene expression levels in bulk cell analyses^11–13^, the relationship between genetic and transcriptional heterogeneity at single-cell levels remains unclear. The rapid development of single-cell RNA and DNA sequencing technologies^14–18^ has allowed researchers to study the transcriptional and genetic heterogeneity of various cancer types at single-cell level. However, simultaneous assessment of RNA and DNA information from the same cell remains technically challenging, limiting studies^19, 20^ that integrate subpopulations characterized by transcript abundance and genetic subclones at single-cell level resolution.

To address this challenge, several methods have been developed recently to infer CNVs from single-cell RNA sequencing (scRNA-seq) data: HoneyBADGER^21^, inferCNV^22^, sciCNV^23^, CaSpER^24^, and CopyKAT^25^. These methods enable the integration analysis of RNA and DNA at the single-cell level. HoneyBADGER^21^ utilizes a hidden Markov model (HMM) integrated with a Bayesian method for CNV detection, providing both expression-based and allele-based CNV inference. Similar to the expression-based method like HoneyBADGER^21^, inferCNV^22^ implements a similar strategy for CNV detection but uses different data de-noising, smoothing, and normalization strategies and considers more CNV states in a transition matrix for CNV detection. sciCNV^23^ calculates an expression disparity score combined with another score that assesses concordant expression changes in contiguous genes to identify CNV. CaSpER^24^ adopts a signal processing approach, performing multiscale smoothing of both gene expression data and allele frequency data and integrating these two types of data for CNV detection. CopyKAT^25^ uses a statistical model to identify and characterize cellular subpopulations, while considering technical noise and biological variation in gene expression for assessing cell-to-cell variation. All five methods claimed to be able to detect both large-scale and focal CNVs. However, these methods have not been evaluated comprehensively with different scRNA-seq platforms or protocols under different sequencing settings and clinical applications.

Here we carried out a comprehensive study to assess the effects of scRNA-seq platform, sequencing depth, and bioinformatics algorithm on scRNA-seq CNV analysis. First, we assessed the sensitivity and specificity of the five CNV inference methods using scRNA-seq data derived from a multicenter benchmarking study on the cell lines (HCC1395/HCC1395BL) based on different scRNA-seq platforms and sequencing depths. We evaluated various factors (including cell numbers, read lengths, read depths, and reference data) for the four scRNA-seq platforms. Second, we evaluated the accuracy of tumor subpopulation identification of each scCNV inference method using a scRNA-seq dataset derived from mixed samples consisting of five human lung adenocarcinoma cell lines. The accuracy was assessed by comparing estimated tumor subpopulations and known cell lines using metrics such as Adjusted Rand Index (ARI)^26^, Fowlkes-Mallows index (FM)^27^, Normalized Mutual Information (NMI)^28^, and V-Measure^29^. Finally, we generated a clinical dataset to validate our findings. Overall, we found that the five methods exhibited large discordance. CaSpER and CopyKAT showed better performance in terms of sensitivity and specificity in CNV inference using scRNA-seq data, whereas inferCNV and CopyKAT exhibited the best performance for identifying tumor subpopulation in a single scRNA-seq platform. However, the expression-based CNV inference methods such as inferCNV, CaSpER, sciCNV, and CopyKAT were highly affected by batch effects (scRNA-seq platform) when estimating tumor subpopulations across multiple platforms. Our evaluation, based on the clinical dataset, showed that CaSpER and CopyKAT outperformed other methods in terms of sensitivity and specificity of CNV inference. However, as regarding subpopulation identification, inferCNV and CopyKAT achieved more accurate results. Our findings revealed the limitations of the current CNV inference methods using scRNA-seq data and provided some guidelines for developing new ones.

## Results

### 1. Overall study design and associated datasets

To evaluate the sensitivity and specificity of the CNV inference methods, we utilized scRNA-seq data derived from our previous multi-center benchmarking study which used paired tumor/normal samples from a human breast cancer cell line (HCC1395, referred to as Sample A) and a matched “normal” control cell line derived from B lymphocytes (HCC1395BL, referred to as B) from the same donor^30–34^. Four scRNA-seq platforms were employed^32^, including two full-length transcript techniques: Fluidigm C1 (referred to as C1) and Takara Bio’s ICELL8 (referred to as ICELL8), as well as two tag-based 3’-transcript techniques: 10x Genomics (hereafter referred to as 10x) and Fluidigm C1 HT (hereafter referred to as C1 HT) (**Figure 1a - Top**). In our study, we first determined the CNVs for the breast cancer cell line based on the bulk cell whole-genome sequencing (WGS) and we detected numerous CNVs^33, 34^. However, to increase the stringency and to make a conservative choice about which of these are most likely true positive CNV events, we chose only 79 CNVs which were also reported by Zack et al.^7^ to be highly recurrent in breast cancers (e.g., the amplification of the MYC oncogene at 8q24.21) as ground truth for our evaluation study. Of these 79 CNVs, 26 CNVs which were identified in our breast cancer cell line using subHMM^35^, based on bulk cell WGS were regarded as true positive events, and the remaining 53 CNVs were considered negative events.

**Figure 1.**
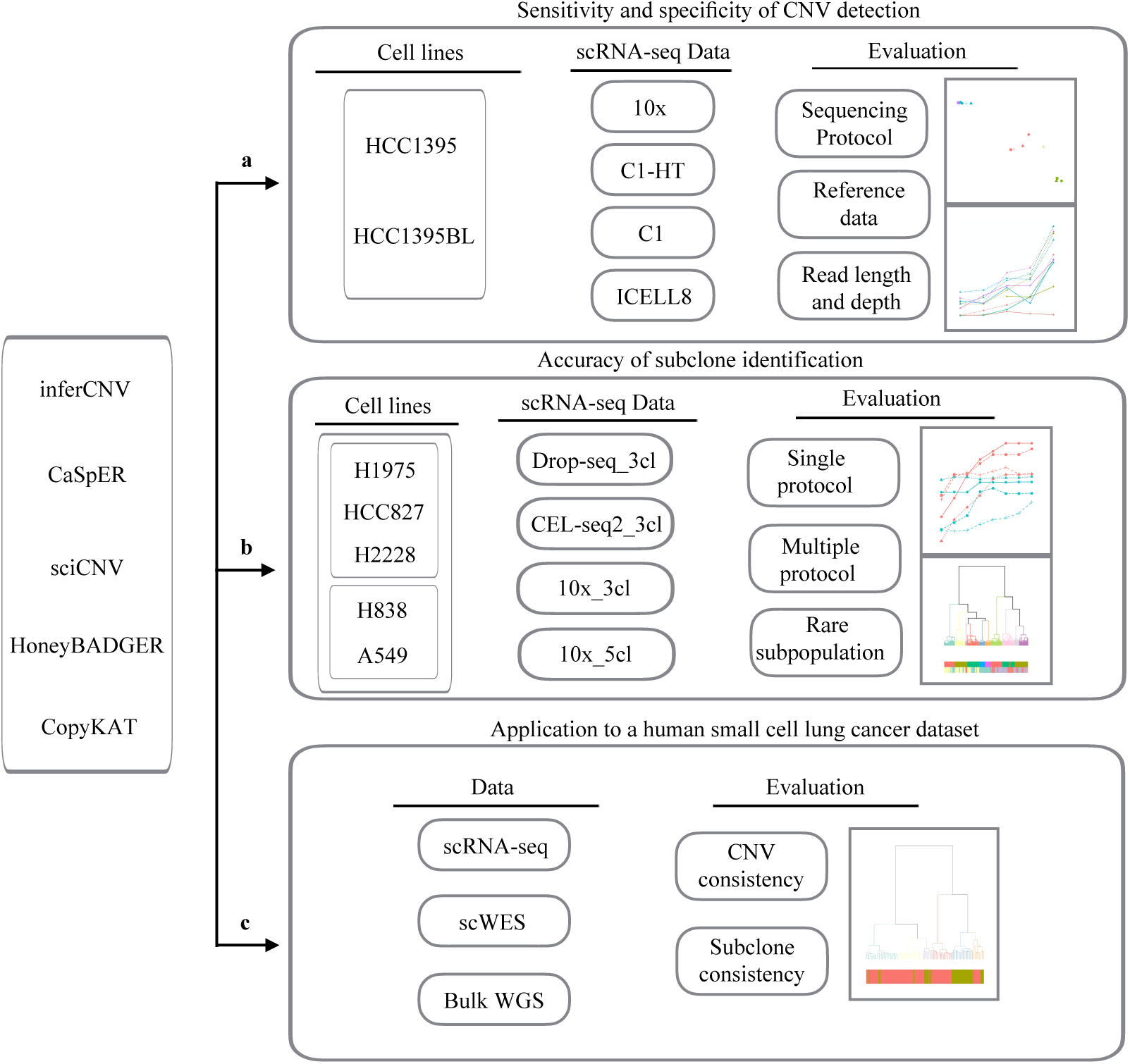
Study design of the scCNV benchmark analysis. The **top panel (a)** illustrates the evaluation scheme for sensitivity and specificity of scCNV detection using the scRNA-seq datasets (10x, C1-HT, C1 and ICELL8 full-length) of a breast cancer cell line vs. the paired B-cell line derived from the donor, which was generated from our previous multicenter benchmarking study. The **middle panel (b)** illustrates the evaluation scheme for accuracy of subclone identification using the mixed scRNA-seq data from Tian et al. study derived from a mixture including either three or five mixed human lung adenocarcinoma cell lines. Drop-seq_3cl, scRNA-seq data from the mixed three human lung adenocarcinoma cell lines, CEL-seq2_3cl, scRNA-seq data from the mixed three human lung adenocarcinoma cell lines. 10x_3cl, 10x scRNA-seq data from the mixed three human lung adenocarcinoma cell lines; 10x_5cl, 10x scRNA-seq data from the mixed five human lung adenocarcinoma cell lines. The **lower panel (c)** illustrates the application of scCNV methods to a human small cell lung cancer (SCLC) scRNA-seq dataset (20M read/each cell, full-length transcript, SMART-seq2) including 92 primary SCLC single cells and 39 relapse SCLC single cells, plus scWES and bulk cell WGS from primary SCLC and relapse tumoral tissues as well as peri-tumoral normal tissues.

To examine the accuracy of tumor subpopulation identification, we used Tian et al. mixed scRNA-seq dataset^36^, derived from mixed samples consisting of either three or five human lung adenocarcinoma cell lines to mimic tumor subpopulations^36^. These datasets were generated using three tag-based techniques: 10x, CEL-seq2, and Drop-seq (**Figure 1b - Middle**).

To evaluate the clinical relevancy and to validate our results based on the above cell line data, we generated a clinical dataset derived from a human small cell lung cancer (SCLC) which is comprised of scRNA-seq data from 92 primary SCLC single cells (20M reads/cell, full-length transcript, SMART-seq2) and 39 relapse SCLC single cells (20M reads/cell, full-length transcript, SMART-seq2), as well as single-cell whole-exome sequencing (scWES, 200x coverage/cell) from 69 primary SCLC single cells, 76 relapse SCLC single cells, and 10 peri-tumoral normal lung single cells plus bulk cell whole-genome sequencing (WGS, 60x coverage/each) data from the primary tumor tissue, relapse tumor tissue and peritumoral tissue (**Figure 1c - Bottom**).

### 2. Sensitivity and specificity of CNV inference by scCNV methods

Cytoband-based CNV comparison was used for the evaluation (see “**Methods”**). Briefly, 79 highly recurrent CNVs^7^, as described above, were considered as total events. These included 34 CNV gains and 45 losses that were denoted by their cytoband locations. CNVs (at cytoband levels) identified by subHMM^35^ in the WGS data were used as ground truth to assess sensitivity and specificity of scCNV callers using scRNA-seq data. Among these 79 highly recurrent CNVs, 26 CNVs (11 CNV gains and 15 CNV losses) identified in the WGS data were considered positive events, while the remaining 53 CNVs were considered negative events. In this study, five scCNV methods, i.e., inferCNV, CaSpER, CopyKAT, sciCNV, and HoneyBADGER (both expression-and allele-based) were applied to scRNA-seq datasets derived from four different scRNA-seq protocols/platforms (**Figure 1a**) to identify CNVs and further annotated at cytoband levels (see “**Methods”**). In addition, we also applied these five scCNV methods to bulk RNA-seq data but found that only CaSpER and CopyKAT could identify CNVs successfully. We included the results of bulk RNA-seq of these two methods in the evaluation.

Overall, CaSpER and CopyKAT showed better performance as compared with the other scCNV methods (**Figure 2a**). These two methods did not show significant performance differences across the four scRNA-seq protocols. To compare the performance of scRNA-seq data derived from four different protocols in more detail, we generated Receiver Operating Characteristic (ROC) curves based on the consensus CNVs identified using different cell percentages (see “**Methods**”, **Figure 2h**). The results are consistent with the above findings, with CaSpER performing slightly better in Fluidigm C1 data, while both methods exhibited less favorable performance in Fluidigm C1-HT data. We extended the evaluation to CNV gains and losses separately (**Figure 2b,c**) and observed differences in performance between CaSpER and CopyKAT. In CNV gains, CopyKAT significantly increased sensitivity but slightly sacrificed specificity compared to CaSpER, except in the Fluidigm C1-HT data (**Figure 2b**). In CNV loss, the performance of the two methods was protocol/scRNA-seq data dependent. CaSpER outperformed CopyKAT in both sensitivity and specificity in Fluidigm C1 and 10x protocol derived scRNA-seq data, whereas CopyKAT showed better specificity in the ICELL8 scRNA-seq data (**Figure 2c**). In addition, CopyKAT showed better performance in CNV gains than in CNV losses. It outperformed in either sensitivity, specificity, or both in the four different protocol derived scRNA-seq data (**Figure 2b,c**). Staggered bar charts (**Figure 2d,e**) illustrated the consistency of the 79 highly recurrent CNVs identified across the five different data sets, including the bulk RNA-seq data, for CaSpER and CopyKAT, respectively.

**Figure 2.**
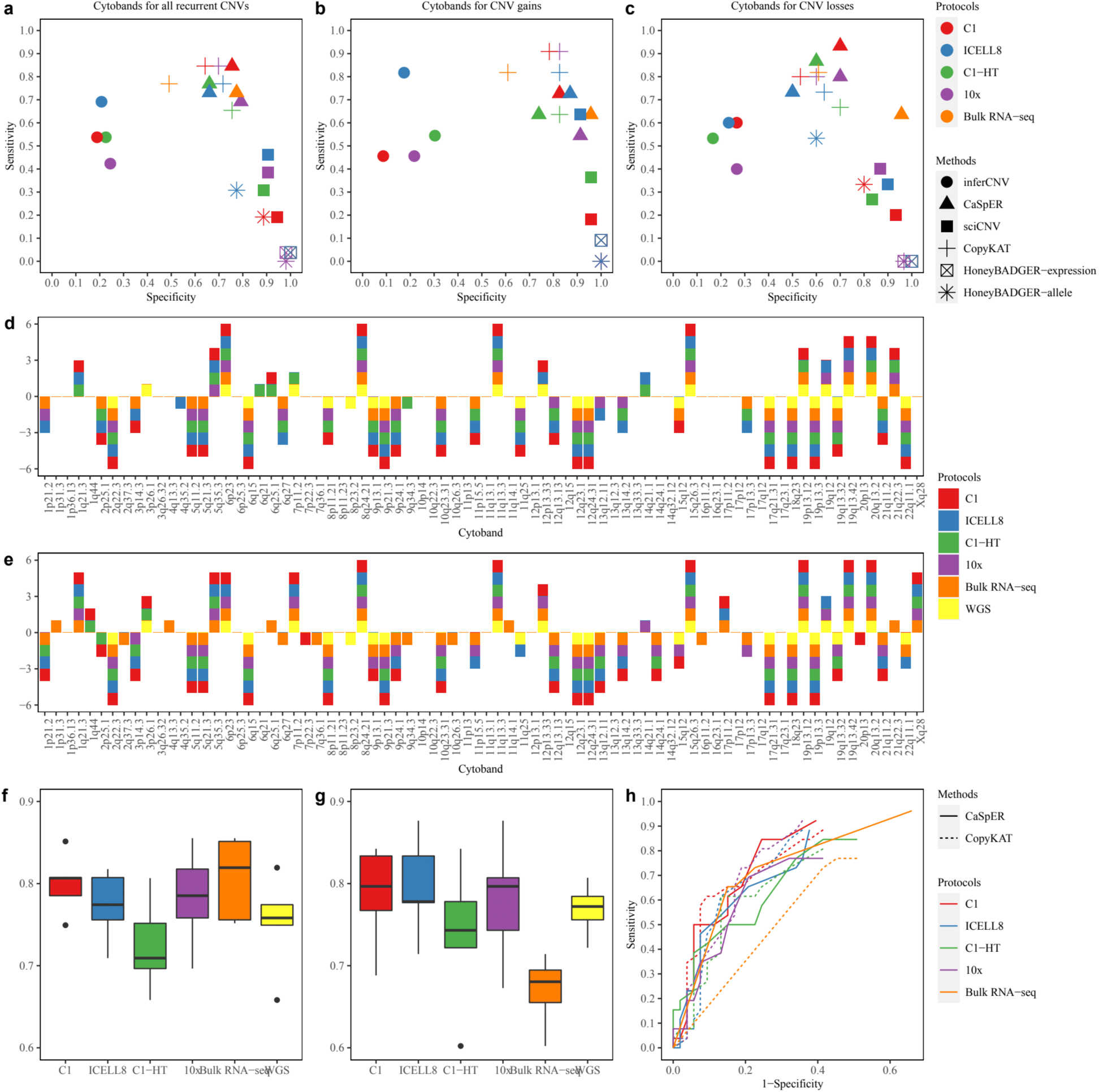
Sensitivity, specificity, and consensus evaluation of scCNV inference methods. **(a-c)** Sensitivity and specificity of scCNV inference methods applied to four different scRNA-seq plus bulk-cell RNA-seq data considering **(a)** all 79 CNVs, **(b)** CNV gains, and **(c)** CNV losses. **(d)** CaSpER and **(e)** CopyKAT showing the CNV identifications based on 79 highly recurrent CNVs in four different scRNA-seq platforms derived datasets along with bulk-cell RNA-seq and bulk-cell WGS dataset. **(f-g)** Consensus of identified CNVs between any two protocols by **(f)** CaSpER and **(g)** CopyKAT. Y-axis refers to ARI score. Each box represents the ARI score between CNV identification status of the x-axis labeled protocol or platform and other protocols or platforms. **(h)** ROC curves of CaSpER and CopyKAT applied to four different scRNA-seq protocols or platforms along with bulk-cell RNA-seq dataset.

In **Figure 2d** and **2e**, consensus CNVs were defined as those identified in all five datasets, while low-concordance CNVs were identified in two or fewer protocol derived data. CopyKAT detected more low-concordance CNVs than CaSpER in both CNV gains (9 vs. 6) and CNV losses (12 vs. 4). Most of these low-concordance CNVs (5 in CNV gains, 7 in CNV losses) originated from the bulk RNA-seq data, indicating a larger number of falsely identified CNVs which contributed to the lower specificity compared with CaSpER (**Figure 2b,c**). In terms of consensus CNVs, CopyKAT detected more CNV gains (9 vs 4), but the number was comparable for CNV losses (14 vs 13) compared with CaSpER, resulting in higher sensitivity for CNV gains across most scRNA-seq protocol derived data.

To assess the consistency of scCNV identification across different scRNA-seq protocol derived data, we measured the similarity of CNV detection status using the Adjusted Rand Index (ARI). Box plots of ARI between a selected scRNA-seq protocol derived data and other scRNA-seq protocol derived data were generated for CaSpER (**Figure 2f**) and CopyKAT (**Figure 2g**). The bulk RNA-seq was also included for ARI analysis. The Fluidigm C1-HT and bulk RNA-seq protocols showed the lowest consistency for CaSpER and CopyKAT, respectively (**Figure 2f,g, Suppl. Figure 1a-d**), suggesting that these methods may not perform optimally for Fluidigm C1-HT and bulk RNA-seq protocol-derived data. The ROC curves (**Figure 2h**) confirmed the subpar performance of Fluidigm C1-HT and bulk RNA-seq with CaSpER and CopyKAT. Furthermore, we evaluated the consistency between any two protocols for CNV gains and losses. In CaSpER, no significant difference was observed between CNV gains and CNV losses (**Suppl. Figure 1e**). In CopyKAT, the ARI score was higher for CNV gains (**Figure S1f**), indicating better performance in detecting CNV gains.

### 3. Effect of scRNA-seq CNV detection using different RNA-seq reference dataset

Reference data as a normal control is required to perform scCNV inference for CNV calling. Depending on the availability of matched normal cells, the scCNV inference methods allow other reference data to perform the analysis. In our evaluation, we examined the impact of three different reference datasets for the four scCNV inference methods, excluding HoneyBADGER due to its extremely low sensitivity.

For the evaluation, we used the following three reference datasets: scRNA-seq data from sample B (HCC1395BL), bulk RNA-seq data from sample B (HCC1395BL), and bulk RNA-seq data of normal breast tissue from the GTex database^37^. Twelve cases in total were evaluated for each scCNV method when three reference datasets and four scRNA-seq protocols were considered. In general, sensitivity decreased significantly in all scCNV inference methods (except inferCNV) across four scRNA-seq protocols when the two bulk RNA-seq reference data were used (**Figure 3a**). An increase in specificity was observed. However, this increase in specificity was not as prominent as the decrease in sensitivity (**Figure 3a**). Noticeably, when using bulk RNA-seq data from GTex database, most scCNV methods showed significantly lower sensitivity and/or specificity (**Figure 3a**).

**Figure 3.**
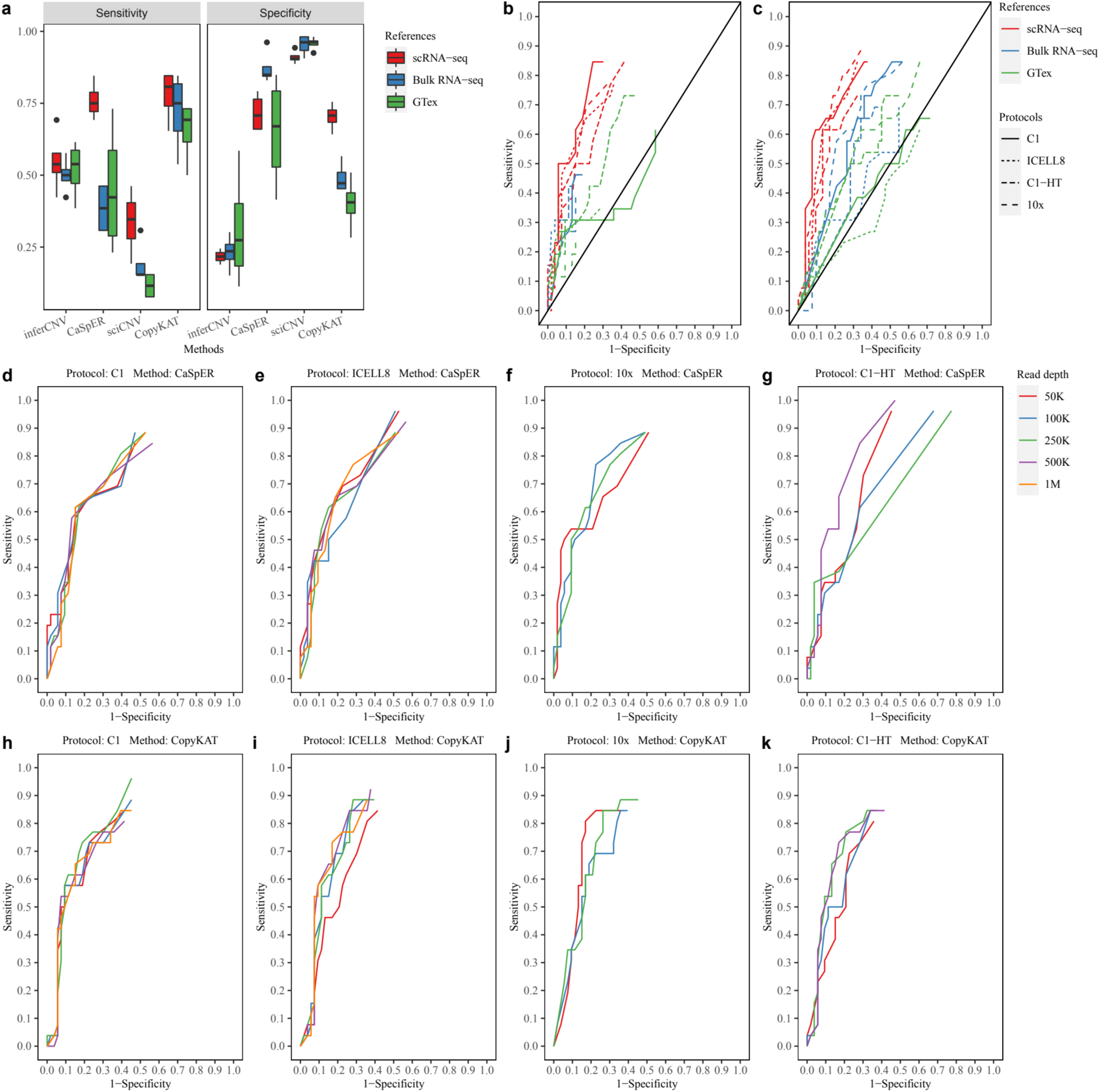
Sensitivity and specificity of scCNV inference methods under multiple conditions. **(a)** Sensitivity and specificity of the four scCNV inference methods by using three reference datasets. Each box represents the sensitivity or specificity of the x-axis labeled method applied to four scRNA-seq protocols. **(b-c)** ROC curves of CaSpER **(b)** and CopyKAT **(c)** applied to four scRNA-seq protocols using three different reference datasets. **(d-k)** ROC curves of CaSpER **(d-g)** and CopyKAT **(h-k)** applied to four scRNA-seq protocols. The reads of each protocol were down sampled to specific read depths.

We analyzed the performance of CaSpER and CopyKAT with the three reference datasets in more detail. The ROC curves of CaSpER (**Figure 3b**) and CopyKAT (**Figure 3c**) suggested that both methods performed better with scRNA-seq reference data as compared to bulk RNA-seq reference data. However, when using the two bulk RNA-seq reference datasets (**Figure 3b**), CaSpER showed low sensitivity in the ROC curves in 6 out of 8 cases because CaSpER failed to call concordant CNVs with high cell percentages (**Suppl. Figure 2**). Among the above 6 cases, more than 57% of CNVs could be detected in fewer than 5% of cells (**Suppl. Table 1**). On the contrary, when using CopyKAT, fewer than 21% of CNVs were detected in fewer than 5% of cells in the 8 worst cases (**Suppl. Table 1**). Our findings indicate that CopyKAT outperformed CaSpER when employing the two bulk RNA-seq reference datasets. Additionally, the ROC curves demonstrated that CopyKAT was more favorable for tag-based scRNA-seq protocols, particularly for 10x protocol, when using the two bulk RNA-seq reference datasets (**Figure 3c**).

### 4. The effect of read length and read depth of scRNA-seq on scCNV inferences

To assess the impact of read length and read depth, our focus was on CaSpER and CopyKAT. We evaluated five different read lengths (50bp, 75bp, 100bp, 125bp, 150bp) and read depths (50k, 100k, 250k, 500k, 1M). For CopyKAT, the ROC curves did not exhibit significant differences across five read lengths and five read depths (**Figure 3h-k, Suppl. Figure 4**). Conversely, for CaSpER, the ROC curves were significantly affected by both read length and depth in three scRNA-seq protocols, except Fluidigm C1 (**Figure 3d-g, Suppl. Figure 3**). Their corresponding AUC showed drastic decreases at specific combinations of read length and read depth (**Figure 3g, Suppl. Figure 3f-i,k,m,n,q**). However, the ROC curves were not monotonically related to the read depth or read length. Further analysis revealed that the unstable performance was due to the usage of different B-allele frequencies (BAFs) files at different read lengths and depths, leading to significantly different BAF shift signal profiles to call CNVs.

To validate this presumption, we utilized 100bp SE data from the ICELL8 protocol (**Suppl. Figure 3h**). The initial analysis (**Suppl. Figure 3m**) showed high and low AUCs at read depths of 500K and 1M, respectively. We reperformed CaSpER analysis on 100bp SE data at five different read depths, but instead of using the BAF files corresponding to each read depth, we replaced them with the BAF files from the read depths at 500K and 1M, respectively (**Suppl. Figure 5a**). For simplicity, we named these two BAF files as 500K-BAF and 1M-BAF. Consistently low AUCs were observed across the five read depths when the 1M-BAF file was used. In contrast, the AUCs increased when the 500K-BAF file was used (**Suppl. Figure 5a**). Similar results were obtained when extending the analysis to other cases in ICELL8 (125bp SE: 500K-BAF vs 250K-BAF) and Fluidigm C1-HT (100bp SE: 250K-BAF vs 500K-BAF; 125bp SE: 500K-BAF vs 250K-BAF) protocols (**Suppl. Figure 5b-d**). Overall, CaSpER was indeed affected by both read length and read depth when using different BAF files.

### 5. Accuracy of scCNV inferred subpopulations using mixed scRNA-seq lung cancer cell line datasets

One of the major applications of scRNA-seq CNV inference methods is the identification of tumor subpopulations. To mimic these subpopulations, we used a mixed sample scRNA-seq dataset GSE118767^36^, which consists of four datasets with mixtures of either three (H1975, H2228, HCC827) or five (A549, H1975, H2228, H838, HCC827) human lung adenocarcinoma cancer cell lines. The mixed scRNA-seq datasets were generated using three different protocols: Drop-seq, CEL-Seq2, or 10x Genomics. We referred to the datasets as Drop-seq_3cl, CEL-seq2_3cl, 10x_3cl and 10x_5cl, respectively. In our evaluation, we considered three or five cancer cell lines as the major subpopulations, making the following assumptions: (1) Each cell line comprises relatively homogeneous clonal cells; and (2) Different cell lines exhibit high heterogeneity. We assessed the performance of the scCNV inference methods using hierarchical clustering trees, ARI, FM, NMI, V-Measure, and number of estimated clusters. We considered three evaluation scenarios for the subpopulation identification: based on a single scRNA-seq protocol or platform, based on combined multiple different scRNA-seq protocols or platforms, and rare subpopulation inference.

#### 5.1. Subpopulation inference based on the data from a single scRNA-seq protocol

In this scenario, we evaluated the scCNV inference methods individually on the four datasets. We processed the Drop-seq and CEL-seq2 datasets using umitools (v1.0.0), while both Cell Ranger V2 and V3 were used to process 10x_3cl and 10x_5cl datasets. This resulted in a total of six gene count matrices (referred to as datasets) for evaluation. Methods that achieved higher ARI, FM, and V-Measure scores demonstrated better performance. HoneyBADGER-expression (expression-based method) could not identify CNVs in any of the 10x datasets, so it was not considered further in the evaluation. Similarly, HoneyBADGER-allele (allele-based method) was not considered further in the evaluation for 10x_5cl datasets. We found that inferCNV, sciCNV, and CopyKAT performed better than CaSpER and HoneyBADGER (**Figure 4**). inferCNV outperformed sciCNV and CopyKAT in the 10x_5cl datasets, suggesting its ability to identify cell subpopulations in more complex and heterogeneous environments (**Figure 4**). Another notable finding was that sciCNV was less affected by 10x_3cl datasets processed by the two Cell Ranger pipelines (**Figure 4, Suppl. Figure 6c,d**). The metrics scores of the two 10x_3cl datasets were similar in sciCNV but not in inferCNV or CopyKAT (**Figure 4**). Interestingly, we observed that most of the additional low RNA content cells called by Cell Ranger V3 were HCC827 cells (631 out of 636 cells). In the case of inferCNV and CopyKAT, hierarchical clustering failed to merge the HCC827 cells called by Cell Ranger V2 and V3 (**Suppl. Figure 6a,b,e,f**). Instead, these methods treated the low RNA content HCC827 cells as a unique subpopulation. To assess the ambient RNA contamination levels in 10x_3cl and 10x_5cl datasets, we conducted a contamination test^1, 38^ and found that the additional cells called by Cell Ranger V3 introduced elevated contamination (**Suppl. Figure 6g**). These findings indicated that inferCNV and CopyKAT may not be a good option to handle low-quality data, as these methods might be sensitive to gene expression changes in individual cells, leading to false CNV identification. Overall, when used with a single protocol, inferCNV, sciCNV, and CopyKAT performed well to identify cell subpopulations. inferCNV might be the best option when the dataset consists of complex cell populations. sciCNV might be better to handle low-quality data.

**Figure 4.**
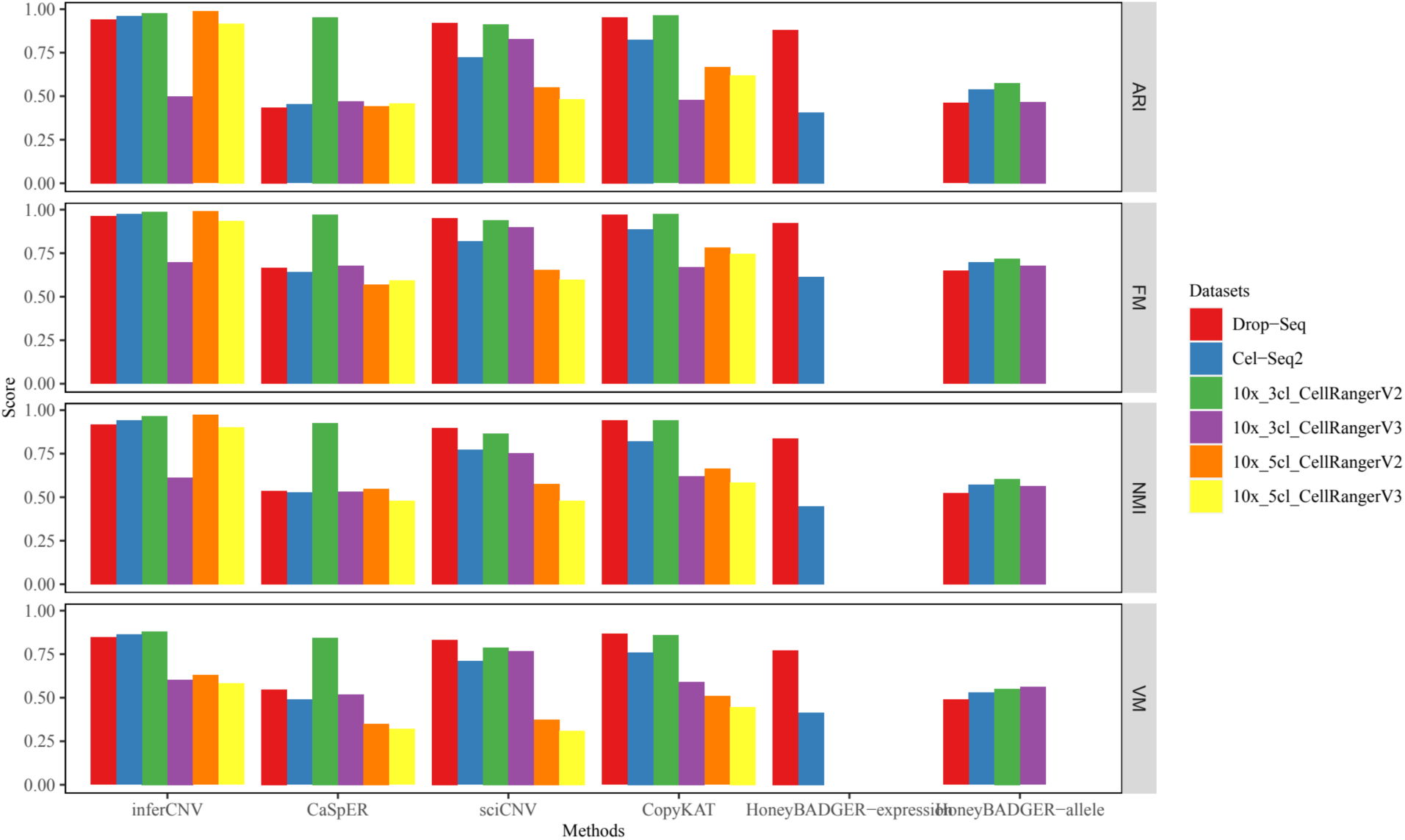
Accuracy of subpopulation identification based on a single scRNA-seq protocol or platform. Four measurement scores **(a)** Adjusted Rand Index (ARI), **(b)** Fowlkes-Mallows index (FM), **(c)** Normalized Mutual Information (NMI), and **(d)** V-Measure were used to evaluate the accuracy in six datasets. The measurement scores were calculated between the estimated clusters and the true cell type labels. The number of clusters estimated was identical to the number of true cell types.

#### 5.2. Subpopulation inference based on combined dataset from multiple scRNA-seq protocols or platforms

Batch effects are a common issue in scRNA-seq analysis. We simulated a test dataset with strong batch effect by combining datasets from different protocols or platforms. In this scenario, we tested the five scCNV inference methods on the combined dataset. To create the combined dataset, we included all cells in the Drop-seq_3cl and CEL-seq2_3cl datasets, and randomly selected 250 cells from each of the 10x_3cl and 10x_5cl (processed by Cell Ranger V2) datasets, resulting in a combined dataset consisting of 943 cells. To evaluate the performance of these methods, we further considered batch based metric scores. Methods that achieved better performance exhibited a strong association between the estimated clusters and the true cell lines, while showing a weak association with batch (representing different scRNA-seq protocols or platforms). The HoneyBAGDER methods were less affected by batch effects, especially for the allele-based method, which achieved the best balance between cell line and batch based metric score when the number of clusters was five (**Figure 5a,p,q**). On the contrary, inferCNV, CaSpER, sciCNV, and CopyKAT were highly affected by batch effects (**Figure 5a,d-g, Figure S5a,d,g**). To improve their performance, we applied batch correction methods before CNV inference. We observed increased cell line-based metric scores and decreased batch based metric scores for all four scCNV inference methods (especially inferCNV and CopyKAT) after applying ComBat batch correction (**Figure 5i,o**). Therefore, for subpopulation inference in the scRNA-seq data combined from multiple scRNA-seq protocols or platforms, inferCNV or CopyKAT combined with ComBat batch correction are the recommended options for inferring cell subpopulations. However, it should be noted that the batch corrected gene expression data may affect the accuracy of detected CNVs. If the batch correction methods are not preferred, the HoneyBADGER methods, especially the allele-based HoneyBADGER, provide the best alternatives.

**Figure 5.**
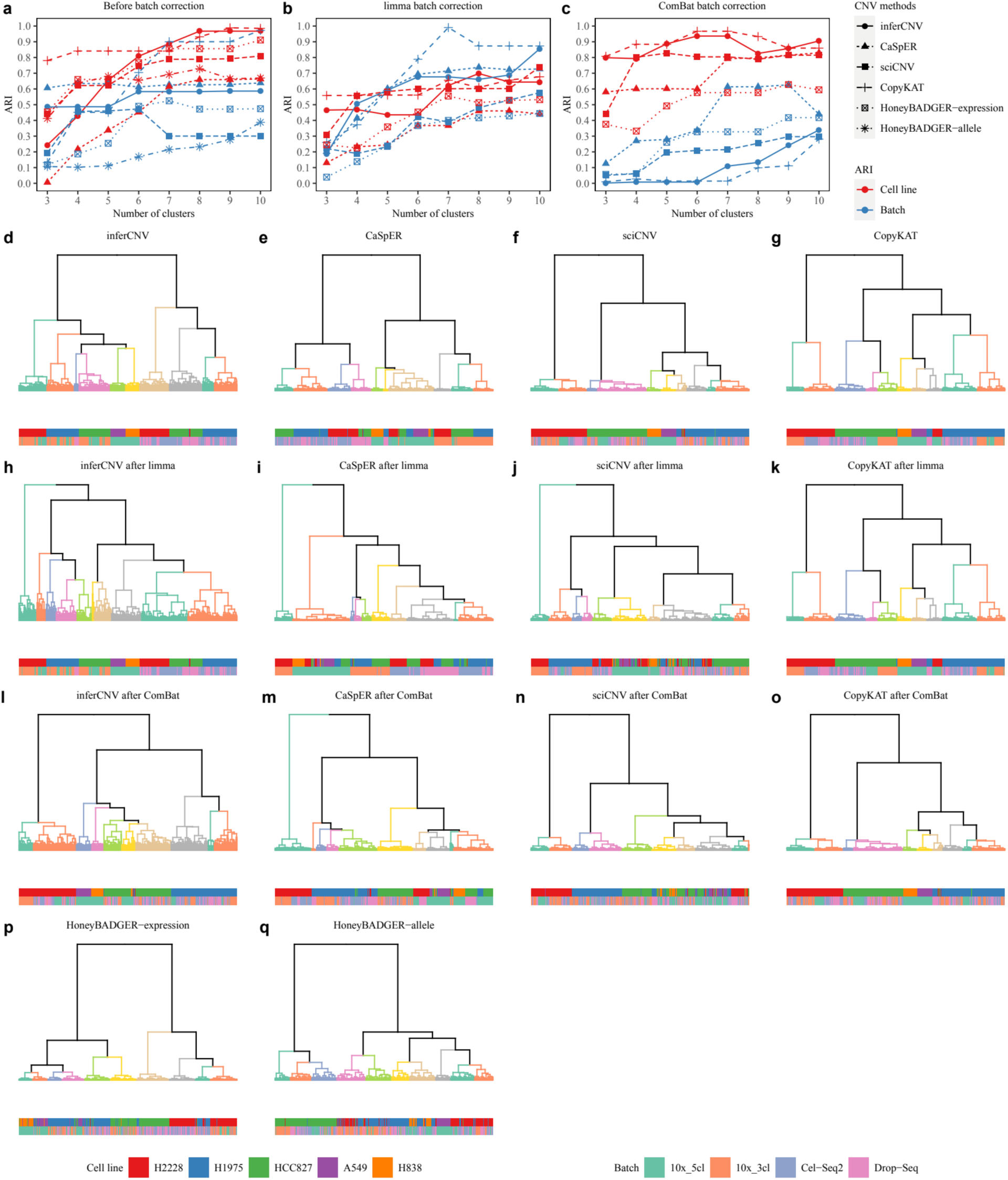
Accuracy of subpopulation identification based on a dataset derived from multiple scRNA-seq protocols or platforms. **(a-c)** The ARI scores between true cell types or batches and the clusters estimated by the five CNV inference methods applied to the dataset **(a)** before, and **(b)** after limma, or **(c)** after ComBat batch corrections. **(d-q)** Hierarchical clustering of the five CNV inference methods applied to the dataset before and after batch correction. The bottom bars represent the true cell line labels and batch labels. HoneyBADGER scCNV method was evaluated at either expression-based or allele-based separately.

#### 5.3. Rare cell subpopulation identification from the mixed lung cancer cell scRNA-seq datasets

In this scenario, we tested the sensitivity of the five methods in identifying rare cell subpopulations using three scRNA-seq datasets from the mixed lung cancer cell lines: Drop-seq_3cl, CEL-seq2_3cl, and 10x_5cl^36^. To simulate a rare cell subpopulation, we selected a low percentage of cells from one specific cell line for evaluation^36^. The experimental design details are provided in **Suppl. Table 4**. In summary, for Drop-seq and CEL-seq2, we randomly selected a total of 100 cells from the three cell lines, with H2228 cells representing the rare cell subpopulation at low percentages (1%, 2%, 5%, and 10%). For 10x_5cl, we considered H1975 cells as the rare cell subpopulation and expanded the design to consider a total of 200, 500, and 1000 randomly selected cells. HoneyBADGER was not evaluated on 10x_5cl data due to its poor performance on the dataset.

Since the scCNV inference methods used Hidden Markov Models (HMM) to infer CNVs, there may be slight variation in the CNVs generated between different runs, which can impact cluster accuracy. To account for this, we repeated each scCNV inference method 10 times for the same dataset. We used three metrics to evaluate the performance: percentage of successful runs, number of clusters to identify the rare cell subpopulation, and proportion of cells labeled as the rare cell subpopulation (see “**Methods**”)

The results of the evaluation are presented in **Suppl. Table 2**. Overall, inferCNV performed best. CopyKAT and CaSpER performed similarly, with CopyKAT slightly outperforming CaSpER in most cases. sciCNV and HoneyBADGER are not recommended for identifying rare cell subpopulations.

We then focused primarily on evaluating inferCNV. For Drop-seq and CEL-seq2, it required the rare population to be at least 5% of cells for the rare subpopulation to be identified. As the cell percentage decreases to 2%, more clusters are required. For 10x, when the total number of cells is small (100 or 200 cells), inferCNV needed at least 5% of cells to be in the rare population to identify the rare subpopulation. However, as the total number of cells increased, inferCNV exhibited increased sensitivity. It could identify the rare subpopulation cells even at 1% when the total number of cells exceeded 500. A similar trend was observed for the other scCNV methods as well, indicating that the sensitivity of the scCNV inference methods in identifying rare subpopulation cells depends on both the percentage and the exact number of rare cells. For inferCNV, if the total number of cells exceeded 500, at least 1% of rare subclonal cells should be detected. If the total number of cells decreased to 200 or lower, a percentage of 5% or higher was required for accurate identification.

### 6. CNV detection using scRNA-seq data derived from a clinical SCLC study

To further validate our findings, we conducted a comparison of CNVs detected by four methods (excluding HoneyBADGER) using a clinical dataset of human small cell lung cancer (SCLC). Similarly, we used CNVs identified by subHMM in WGS data from primary and relapse samples of the same patient as the ground truth. For the comparison of cytoband-based CNVs, we considered 54 highly recurrent CNVs (denoted as cytobands) from a reference paper^7, 39^ as the total events. These cytobands could be divided into 27 CNV gains and 27 CNV losses. In the primary scRNA-seq SCLC data, we observed similar results to our above evaluation (**Figure 6a**). CaSpER and CopyKAT exhibited the best performance in terms of sensitivity and specificity. However, in the relapse data, these two scCNV inference methods showed low specificity. This discrepancy might be attributed to the selection of the 54 highly recurrent CNVs, which were mainly detected in primary SCLC samples. In the evaluation of subpopulation, the results were consistent with the conclusions drawn above (see **Section 5.1**). inferCNV and CopyKAT outperformed the other methods and were able to cluster primary and relapse cells with high accuracy **(Figure 6b-e)**.

**Figure 6.**
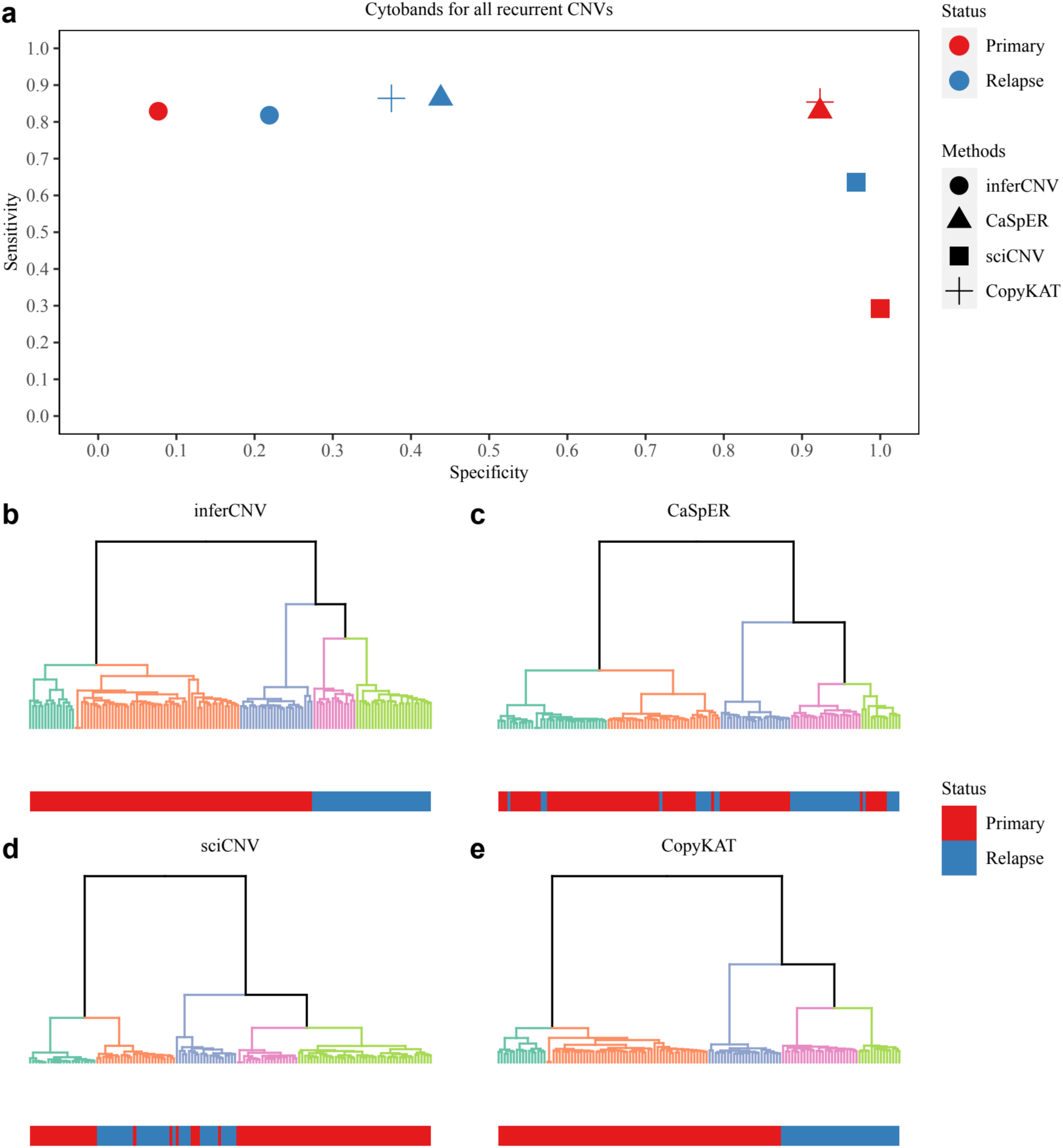
Evaluation of scCNV inference methods using a clinical small cell lung cancer (SCLC) dataset. **(a)** Sensitivity and specificity of the four scCNV inference methods applied to primary and relapse SCLC dataset. **(b-e)** Hierarchical clustering of the four scCNV inference methods applied to the SCLC dataset.

## Discussion

In this study, we evaluated the capability of five scCNV inference methods across 8 datasets representing 6 different scRNA-seq protocols or platforms including both tag-based and full-length transcript scRNA-seq technologies^32, 40^. We also applied the methods to a clinical dataset for validation. Two major aspects were considered in the evaluation, accuracy (sensitivity and specificity) of inferred CNVs and accuracy of identified subclones. To evaluate the accuracy of inferred CNVs, the CNVs identified by WGS data were used as ground truth^7, 33, 34^. In the evaluation, four different factors were considered: number of cells, normal reference samples, read length, and sequencing depth. We observed large variations of CNVs inferred by the five scCNV inference methods, resulting in quite different sensitivity and specificity (**Figure 2a-c**). HoneyBADGER was the most conservative method; it identified the fewest CNVs and had the lowest sensitivity and highest specificity. inferCNV and sciCNV showed opposite patterns in terms of sensitivity and specificity, indicating either high false negative or low true positive rate. CaSpER and CopyKAT achieved the best balance between sensitivity and specificity.

The selection of normal reference datasets showed different effects on scCNV inference methods. Overall, best performance can be achieved when using scRNA-seq reference data (**Figure 3a**). When the two bulk RNA-seq reference datasets were used, CopyKAT outperformed other methods (**Figure 3a-c**). Furthermore, we confirmed that CopyKAT worked extremely well for tag-based scRNA-seq protocols or platforms, especially the 10x platform, with all three normal reference datasets (**Figure 3c**).

When evaluating the effect of read length and read depth, an interesting finding was that CaSpER led to inconsistent sensitivity and specificity at different read depths of the same sample (**Figure 3g, Suppl. Figure 3**). We believe the inconsistency is associated with a large variation of BAF shift thresholds estimated from different BAF signal profiles at each read depth. For example, in the ICELL8 125bp case, we found the highest BAF shift thresholds at a read depth of 250K as compared with the other four read depths (**Suppl. Table 3**). In the CaSpER algorithm, the estimation of BAF shift threshold is critical for CNV filtering. The high BAF shift threshold resulted in a large number of filtered CNVs and low sensitivity. In our analysis, we used only the recommended parameter settings to extract BAF. A better fine-tuning of parameters in the current BAF extracting algorithm or developing a better BAF extracting algorithm might improve CaSpER analysis substantially.

In terms of subpopulation identification, inferCNV and CopyKAT showed better performance in the single protocol derived scRNA-seq data compared to the multiple protocols derived scRNA-seq data using the mixed samples consisting of either three or five different lung cancer cell lines^36^. However, in multiple protocol inference, all methods except HoneyBADGER were deeply affected by batch effects^32^. We tested two batch correction methods, i.e., limma^41^ and ComBat^42^ designed for bulk RNA-seq and found ComBat could improve the performance of inferCNV and CopyKAT dramatically. The recently developed scRNA-seq batch correction methods seem not to work well with current scRNA-seq CNV inference methods because most of them only provided embedding matrices or normalized gene expression matrices instead of batch corrected gene expression matrices^32^. Since batch effects strongly affected the scRNA-seq CNV inference methods, we believe it is essential to develop novel scRNA-seq batch correction methods that overcome this limitation.

Computational runtime is critical for single cell analysis because current scRNA-seq techniques can generate gene expression data with from a few hundred to hundreds of thousands of single cells. In our study, we tested the dataset used in the “multiple protocol inference” section with a total of 943 cells and further down sampled to 100, 200, and 500 cells. Since inferCNV supports multicore running, we evaluated different numbers of cores (1, 5, 10, and 20) to run inferCNV. All runs were conducted on a server with 2x Intel Xeon E7-8860 CPUs at 2.2 GHz, 384 GB memory. CopyKAT required the least computational time. HoneyBADGER-allele based method requires an excessively long time (> 24 h) when 500 or more cells are analyzed (**Suppl. Figure 7**). CaSpER and HoneyBADGER-expression based methods required less time compared with inferCNV with less than 20 cores. inferCNV benefits a lot from multicore running, especially when the number of cells increases. With 943 cells, the three methods CaSpER, HoneyBADGER-expression, and inferCNV (with 20 cores) had similar computational time (∼ 2 h).

## Conclusions

We observed large differences in performance between the five scCNV inference methods evaluated. CaSpER and CopyKAT showed better sensitivity and specificity, while inferCNV and CopyKAT were best for identifying tumor subpopulations using a “single protocol or platform” derived scRNA-seq data as compared to the “multiple protocols or platforms” derived scRNA-seq data in the mixture scRNA-seq datasets^36^. However, the expression based scCNV inference methods such as inferCNV, CaSpER, sciCNV, and CopyKAT were highly affected by batch effects when estimating tumor subpopulations in mixed samples using multiple protocols. In the clinical application, CaSpER and CopyKAT outperformed other methods in terms of sensitivity and specificity. Regarding subpopulation identification, inferCNV and CopyKAT achieved better performance. Our findings address the limitations of the current methods and provide guidelines for developing new ones.

## Methods

### Datasets used in our study

We used three different data sets in our scCNV inference method study including scRNA-seq data from a multicenter benchmark study^32^, scRNA-seq data from mixed human lung adenocarcinoma cell line samples^36^ and the newly generated clinical human small cell lung cancer data. These datasets were briefly described here as follows.

### scRNA-seq datasets from a multicenter study

We used the scRNA-seq datasets from our previous multi-center benchmarking study which were obtained from a human breast cancer cell line (HCC1395) vs. the matched “normal” control B lymphocyte cell line (HCC1395BL) from the same donor^30–34^. The scRNA-seq datasets were generated from four scRNA-seq platforms including two full-length transcript techniques: Fluidigm C1 (referred to as C1) and Takara Bio’s ICELL8 (referred to as ICELL8), and two tag-based 3’-transcript techniques: 10x Genomics (hereafter referred to as 10x) and Fluidigm C1 HT (hereafter referred to as C1 HT). The detailed methods on how the scRNA-seq data were generated were described in our previous publication^32^. The multi-center scRNA-seq fastq raw data is available in the SRA repository with the access code #: PRJNA504037.

### scRNA-seq data from mixed human lung adenocarcinoma cell line samples

We also used Tian et al. mixed scRNA-seq dataset^36^, derived from mixed samples consisting of either three or five human lung adenocarcinoma cell lines to mimic tumor subpopulations^36^. These datasets were generated using three tag-based techniques: 10x, CEL-seq2, and Drop-seq. The detailed methods on how these mixed scRNA-seq data were generated can be found in Tian et al paper^36^. The fastq raw data from Tian et al. mixture scRNA-seq data for the five lung cancer cell lines are available under GEO SuperSeries GSE118767.

### Clinical human small cell lung cancer data

#### Tissue collection

All sample collection, patient consent, and patient recruitment followed Institutional Review Board (IRB) protocol approved by the Hangzhou Cancer Hospital, Hangzhou, China. The samples were obtained from a 59-year-old male with an initial diagnosis of left lung cancer by a chest computed scan. The patient further underwent CT-guided lung biopsy for pathology examination which confirmed a diagnosis of extensive stage small cell lung cancer (SCLC) with liver and bone metastasis. The patient was subject to chemotherapy. Before the chemotherapy, we obtained three pieces of cancer tissues (0.2×1.5 cm/each) by CT-guided lung puncture which were further subject to hematoxylin and eosin (HE) staining for confirmation. A 10 ml blood sample was obtained as control. One piece of cancer tissue sample was stored in −80 °C, and other two pieces of tissue samples were immediately used for single-cell isolations. When the patient was recurred after chemotherapy, two pieces of cancer tissues (0.2×1.5 cm/each) were obtained by CT-guided lung puncture. One piece of cancer tissue was stored in −80 °C, and two other pieces of tissue samples were used for single-cell isolations. For the tissue collections both during the primary and relapse periods, adjacent normal tissues (3 perforated tissues) were also obtained from the patient which were subject to HE stains and pathology confirmation as no cancer tissues (peritumoral normal tissues).

#### Bulk cell WGS

genomic DNA (gDNA) was extracted from the fresh frozen tissues using Qiagen QIAamp DNA mini kit (Qiagen, Germany) according to manufacturer’s protocol. The DNA sample quality and integrity were evaluated by A260/A280 ratio and agarose gel electrophoresis. The concentration of gDNA was determined by Nanodrop 2000 (Thermo, USA) and Qubit 3.0 (Life Technologies, USA). WGS libraries were constructed using Illumina Tru-seq Nano DNA HT Sample Prep kit with 0.5 µg gDNA following the manufacturer’s protocol and sequenced on an Illumina X10, paired end (PE, 150 bpx2) with 30x coverage. The fastq sequencing data were deposited in the SRA database, under SRA number SRP149859.

#### Single cell isolation

The tissues were digested with collagenase IV (Sigma Aldrich, Shanghai, China), at 37 °C for 30 min, washed in phosphate-buffered saline and the cells were then diluted to 1000 cells/ml. Single cells were isolated using a micromanipulation system^43^.

#### Single-cell RNA-seq (scRNA-seq)

cDNAs were amplified from 92 primary tumor cells and 39 relapse tumor cells using SMART-seq2 kit as described previously^44^. scRNA-seq libraries were constructed with 1 ng amplified cDNA using Nextera XT DNA Library Preparation kit (Illumina, San Diego, CA, USA) following the manufacturer’s protocol. The scRNA-seq libraries were sequenced with 20M reads per cell on an Illumina X10 with PE reads, 150 pbx2. The fastq sequencing data were deposited in the SRA database, under SRA number SRP149859.

#### Single-cell WES (scWES)

Single-cell genomic DNA obtained from 69 primary tumor cells and 76 relapse tumor cells were amplified with Qiagen REPLI-g Single Cell kit (Qiagen, Shanghai, China) as described previously^43^. Exome capture was performed on amplified DNA from each single cell using the Agilent SureSelect Clinical Research Exome panel (Agilent Technologies, Shanghai, China) following the manufacturer’s protocol. The Agilent SureSelect Clinical Research Exome panel targets a 54 Mb region including gene exons. The captured DNAs were purified using the AMPure XP beads (Beckman Coulter). The scWES libraries were sequenced on an Illumina X10 with PE reads, 150 bpx2, with 200x coverage for each cell. The fastq sequencing data was deposited in the SRA database, under SRA number SRP152993.

### Human reference genome

The reference genome and transcriptome were downloaded from the 10X website as refdata-cellranger-GRCh38-1.2.0.tar.gz. This reference corresponds to the GRCh38 genome and Ensmebl v84 transcriptome. All data analyses were performed using this reference genome and transcriptome.

### Preprocessing of scRNA-seq data

For 10x data, the raw fastq data were processed usingCellRanger (v2.1.0 and v3.1.0) to generate gene count matrices. In the CellRanger pipeline, cellranger count was used with all default parameter settings.

For C1, C1-HT, and ICELL8 data, the quality of the raw fastq data was assessed by fastqc. cutadapt (v1.9.1) was used for trimming and filtering. Bases with quality less than 10 were trimmed from 5’ and 3’ ends of reads. Reads less than 20 bases were excluded from further analysis. STAR (v2.5.4b) with default parameter settings was used for alignment to generate bam files. The featureCounts (v1.6.1) was used to generate gene counts per cell. All default parameter settings were used in the gene counting.

For Tian’s Drop-seq and CEL-seq2 data, umitools (v1.0.0) was used to process the raw fastq data and generate gene count matrices. In the umitools pipeline, ‘umi_tools whitelist’ with default parameter settings was used to generate a list of cell barcodes for downstream analysis. ‘umi_tools extract’ was used to extract the cell barcodes and filter the reads if phred sequence quality of cell barcode bases was < 10 or UMI bases < 10 (options: --quality-filter-threshold=10 --filter-cell-barcode). STAR was used for alignment to generate bam files containing the unique mapped reads (option: outFilterMultimapNmax 1) for gene counting. featureCounts was used to assign reads to genes and generate a BAM file (option: -R BAM). Finally, ‘umitools count’ (options: --per-gene --gene-tag=XT --per-cell --wide-format-cell-counts) was used for the sorted BAM files to generate gene count per cell matrices.

### CNV inference methods

Heterozygous SNP identification for HoneyBADGER: Three heterozygous SNP reference sets were used in the HoneyBADGER allele-based method. For SNPs sourced from ExAC, common heterozygous variants (in VCF format) were extracted from the database. These variants underwent filtering to include only common SNPs with a minor allele frequency (MAF) greater than 10%.

Consensus SNPs identified from 63 HCC1395BL WGS samples followed the data processing and VCF extraction guidelines outlined in a reference paper^33^. The same variant filtering strategy was applied to obtain the final SNP references.

In one WES sample, the fastq files were trimmed using cutadapt (v1.9.1). Reads less than 20 bases were excluded from further analysis. The trimmed fastq files were aligned to the human reference genome (GRCh38) using BWA MEM (v0.7.13). Duplicates were removed using Picard tools (v1.141). Indel realignment and quality score recalibration were performed using GATK v3.8.1. Germline heterozygous variants were identified using GATK’s HaplotypeCaller, followed by variant quality score recalibration. The same variant filtering strategy was applied to obtain the final SNP reference set.

HoneyBADGER: For expression-based analysis (HoneyBADGER-expression), the log-transformed counts per million (CPM) were used as input, and the following gene filtering parameters were applied: Genes with the mean expression lower than 1 CPM in both test and control samples, or genes with the mean expression lower than 3.5 CPM in the test sample, or genes with the mean expression lower than 2.6 CPM in the control sample were filtered for further analysis.

For the allele-based analysis (HoneyBADGER-allele), SNPs with greater than 0.05 deviation from the expected 0.5 heterozygous allele fraction were filtered out for further analysis.

InferCNV: The inferCNV analysis involved the following steps: “denoise” was performed using the default hidden markov model (HMM) settings, with a “cutoff” value of 0.1 for tag-based protocols and 1 for full-length protocols. The “subcluster” method was applied to infer the subcluster cells using the “random_trees” partition method. The default p-value of 0.05 was used to determine cut-points in the hierarchical tree.

CaSpER: BAFExtract was used to obtain BAF for each bam file of the scRNA-seq data. The log-transformed counts per million (CPM) were used as gene expression matrices. The CaSpER analysis was performed in “single-cell” mode, with expressed genes filtered out when their expression was less than 0.1.

sciCNV: The gene counts matrices were normalized using RTAM2 normalization method developed by the same group. The sciCNV analysis was performed on the normalized gene expression matrices with default parameter settings. The sciCNV scores per gene per cell were calculated for further analysis.

CopyKAT: CopyKAT analysis was performed using default parameter settings.

### Cytoband-based CNV comparison

In the cytoband-based CNV comparison, we started by defining the ground-truth CNVs based on their cytoband locations. This allowed us to generate total events, positive events, and negative events, and further calculated sensitivity and specificity. We considered CNVs reported from the two reference papers^7, 39^ as total events. To determine the positive events, we identified the CNVs that overlapped between the total events and the CNVs identified in the WGS data using subHMM. The CNVs that did not overlap were considered negative events.

Due to the variations in CNV output among the five scCNV inference methods, we converted the CNV output into gene-based CNVs for downstream analysis. This involved assigning CNVs to genes at single cell level if they overlapped with the genomic region of the CNVs. Subsequently, we defined consensus gene-based CNVs if the genes were called in more than 10% of the cells. Using the cytoband location of gene-based CNVs and the ground truth, we calculated sensitivity and specificity. ROC curves were generated based on the consensus gene-based CNVs with different cell percentage cutoffs ranging from 5% to 100%.

### Cell type identification for Tian’s data

To determine the cell types of Tian’s scRNA-seq data, we initially obtained bulk RNA-seq data from the five lung cancer cell lines (GSE64098)^45^. We performed a differential gene expression analysis on this bulk RNA-seq data to identify the top 5 gene marks for each of the five cell lines.

For scRNA-seq datasets, we applied the standard preprocessing and clustering steps using the Seurat package. The identified gene marks for each of the cell lines from the bulk RNA-seq data were used as inputs to generate feature plots to determine the cell lines for estimated clusters.

### Metrics score calculation

To calculate the four metrics scores (ARI, FM, NMI, and V-Measure), we used several R packages: fossil (v0.4.0), dendextend (v1.13.4), aricode (v1.0.0), and infotheo (v1.2.0) Each of these packages was used for specific calculations related to the four metrics. To calculate the scores, hierarchical clustering was performed on the outputs of the five CNV inference methods. Then the metrics scores were calculated based on the cluster labels in hierarchical clustering and the true cell type labels or batch labels. Here is a description of the hierarchical clustering approach used for each CNV inference method:

- HoneyBADGER: hierarchical clustering was applied to the posterior probability of each CNV in each cell.
- inferCNV: tumor subclustering was used to obtain hierarchical clustering results.
- CaSpER: hierarchical clustering was applied to smoothed genome-wide gene expression data.
- sciCNV: hierarchical clustering was applied to sciCNV scores derived from the method.
- CopyKAT: the default hierarchical clustering provided by the CopyKAT method was used.

### Metrics to evaluate rare subpopulation identification

To evaluate the performance of the five scCNV inference methods in identifying rare subpopulations, we employed three metrics: percentage of success run, number of clusters required to identify the rare subpopulation cells (referred to as number of clusters), and the proportion of cells labeled as rare subpopulation cells (referred to as cell proportion).

Hierarchical clustering was applied to the datasets designed to estimate clusters. To determine the number of clusters, we constructed a contingency table comparing the estimated cluster labels with the cell line labels. Initially, we set the estimated number of clusters to three. We gradually increased the number of clusters until we obtained one cluster that included only the rare subpopulation cells. The upper limit of the number of clusters was defined as 30. If the number of clusters exceeded 30, the run was considered a failure.

To ensure robustness, we performed a total of 10 repeated runs for the same dataset using each scCNV inference method, considering the HMM approach employed in these methods. The percentage of successful runs represented the number of successful runs out of 10 repeated runs for each dataset and scCNV method. The cell proportion metrics indicated the proportion of rare subclonal cells within the estimated cluster compared with the total number of rare subclonal cells.

## Data availability

The datasets generated and analyzed (multi-center scRNA-seq data) in the current study are available in the SRA repository with the access code #: PRJNA504037, and the following URL: https://www.ncbi.nlm.nih.gov/bioproject/PRJNA504037

The raw data from Tian et al. mixture scRNA-seq data for the five lung cancer cell lines are available under GEO SuperSeries GSE118767, and the following URL: https://www.ncbi.nlm.nih.gov/geo/query/acc.cgi?acc=GSE118767

The raw data for human SCLC are available in SRA repository with the accession code #: SRP152993 (scWES and bulk cell WGS), SRP149859 (scRNA-seq) and the following URL: https://trace.ncbi.nlm.nih.gov/Traces/?view=study&acc=SRP15299 https://trace.ncbi.nlm.nih.gov/Traces/?view=study&acc=SRP149859

## Code availability

The algorithms and code sets for our bioinformatics analyses have been published previously.

## Acknowledgements

The authors would like to thank Ms. Diana Ho and Ms. Adriana Lopez of the LLU Center for Genomics for their administrative support, particularly in coordinating the Zoom conference calls for the project. The authors would like to thank ATCC, and particularly Liz Kerrigan for providing the two cell lines, i.e., HCC1395 and HCC1395BL for our study. The genomic work carried out at the LLU Center for Genomics was funded in part by the Ardmore Institute of Health grant 2150141 (CW) and Dr. Charles A. Sims’ gift to LLU Center for Genomics. The work of Chunlin Xiao was supported by the National Center for Biotechnology Information of the National Library of Medicine (NLM), National Institutes of Health.

## Authors’ contributions

CW conceived, designed the study, and provided funding. CW and CX managed the project. SW provided the human SCLC clinical data. XC drafted the manuscript, conducted all the bioinformatics data analyses, and generated all the figures and tables. WC, ZC and HW carried out genomics experiments. LTF and BZ helped bioinformatics data analysis. FZ, LS and WX helped with the finalizations of figures, tables, and cited references. CX, WJ, MMJ, AF, XZ, SG, CM, BZ, LTF and CW helped edit the manuscript. All authors reviewed the manuscript. CW revised and finalized the manuscript.

## Reference

1. Rosenthal, R., McGranahan, N., Herrero, J. & Swanton, C. Deciphering Genetic Intratumor Heterogeneity and Its Impact on Cancer Evolution. Annu Rev Canc Biol 1, 223–240 (2017).

2. Marusyk, A., Almendro, V. & Polyak, K. Intra-tumour heterogeneity: a looking glass for cancer? Nat Rev Cancer 12, 323–334 (2012).

3. Andor, N. et al. Pan-cancer analysis of the extent and consequences of intratumor heterogeneity. Nat Med 22, 105–+ (2016).

4. Polyak, K. Heterogeneity in breast cancer. J Clin Invest 121, 3786–3788 (2011).

5. Landau, D.A. et al. Evolution and Impact of Subclonal Mutations in Chronic Lymphocytic Leukemia. Cell 152, 714–726 (2013).

6. Burrell, R.A., McGranahan, N., Bartek, J. & Swanton, C. The causes and consequences of genetic heterogeneity in cancer evolution. Nature 501, 338–345 (2013).

7. Zack, T.I. et al. Pan-cancer patterns of somatic copy number alteration. Nat Genet 45, 1134–U1257 (2013).

8. Meijers-Heijboer, H. et al. Low-penetrance susceptibility to breast cancer due to CHEK2*1100delC in noncarriers of BRCA1 or BRCA2 mutations. Nat Genet 31, 55–59 (2002).

9. Beroukhim, R. et al. The landscape of somatic copy-number alteration across human cancers. Nature 463, 899–905 (2010).

10. Bignell, G.R. et al. Signatures of mutation and selection in the cancer genome. Nature 463, 893–U861 (2010).

11. Stranger, B.E. et al. Relative impact of nucleotide and copy number variation on gene expression phenotypes. Science 315, 848–853 (2007).

12. Fehrmann, R.S.N. et al. Gene expression analysis identifies global gene dosage sensitivity in cancer. Nat Genet 47, 115–125 (2015).

13. Martinez-Climent, J.A. et al. Transformation of follicular lymphoma to diffuse large cell lymphoma is associated with a heterogeneous set of DNA copy number and gene expression alterations. Blood 101, 3109–3117 (2003).

14. Zheng, G.X.Y. et al. Massively parallel digital transcriptional profiling of single cells. Nat Commun 8 (2017).

15. Gao, R.L. et al. Nanogrid single-nucleus RNA sequencing reveals phenotypic diversity in breast cancer. Nat Commun 8 (2017).

16. Macosko, E.Z. et al. Highly Parallel Genome-wide Expression Profiling of Individual Cells Using Nanoliter Droplets. Cell 161, 1202–1214 (2015).

17. Hashimshony, T. et al. CEL-Seq2: sensitive highly-multiplexed single-cell RNA-Seq. Genome Biol 17 (2016).

18. Picelli, S. et al. Full-length RNA-seq from single cells using Smart-seq2. Nat Protoc 9, 171–181 (2014).

19. Macaulay, I.C., et al. G&T-seq: parallel sequencing of single-cell genomes and transcriptomes. Nature Methods 12, 519–+ (2015).

20. Dey, S.S., Kester, L., Spanjaard, B., Bienko, M. & van Oudenaarden, A. Integrated genome and transcriptome sequencing of the same cell. Nat Biotechnol 33, 285–+ (2015).

21. Fan, J. et al. Linking transcriptional and genetic tumor heterogeneity through allele analysis of single-cell RNA-seq data. Genome Res 28, 1217–1227 (2018).

22. Patel, A.P. et al. Single-cell RNA-seq highlights intratumoral heterogeneity in primary glioblastoma. Science (New York, N.Y.) 344, 1396–1401 (2014).

23. Mahdipour-Shirayeh, A., Erdmann, N., Leung-Hagesteijn, C. & Tiedemann, R.E. sciCNV: high-throughput paired profiling of transcriptomes and DNA copy number variations at single-cell resolution. Brief Bioinform 23 (2022).

24. Harmanci, A.S., Harmanci, A.O. & Zhou, X.B. CaSpER identifies and visualizes CNV events by integrative analysis of single-cell or bulk RNA-sequencing data. Nat Commun 11 (2020).

25. Gao, R.L. et al. Delineating copy number and clonal substructure in human tumors from single-cell transcriptomes. Nat Biotechnol 39, 599–608 (2021).

26. Hubert, L. & Arabie, P. Comparing Partitions. J Classif 2, 193–218 (1985).

27. Fowlkes, E.B. & Mallows, C.L. A Method for Comparing Two Hierarchical Clusterings. Journal of the American Statistical Association 78, 553–569 (1983).

28. Viola, P. & Wells, W.M. Alignment by maximization of mutual information. Int J Comput Vision 24, 137–154 (1997).

29. Koehn, P. & Hoang, H. Proceedings of the 2007 Joint Conference on Empirical Methods in Natural Language Processing and Computational Natural Language Learning (EMNLP-CoNLL). The Association for Computational Linguistics (2007).

30. Talsania, K. et al. Structural variant analysis of a cancer reference cell line sample using multiple sequencing technologies. Genome Biol 23, 255 (2022).

31. Xiao, C. et al. Personalized genome assembly for accurate cancer somatic mutation discovery using tumor-normal paired reference samples. Genome Biol 23, 237 (2022).

32. Chen, W. et al. A multicenter study benchmarking single-cell RNA sequencing technologies using reference samples. Nat Biotechnol 39, 1103–1114 (2021).

33. Fang, L.T. et al. Establishing community reference samples, data and call sets for benchmarking cancer mutation detection using whole-genome sequencing. Nat Biotechnol 39, 1151–1160 (2021).

34. Xiao, W. et al. Toward best practice in cancer mutation detection with whole-genome and whole-exome sequencing. Nat Biotechnol 39, 1141–1150 (2021).

35. Choo-Wosoba, H., Albert, P.S. & Zhu, B. A hidden Markov modeling approach for identifying tumor subclones in next-generation sequencing studies. Biostatistics 23, 69–82 (2022).

36. Tian, L.Y. et al. Benchmarking single cell RNA-sequencing analysis pipelines using mixture control experiments. Nature Methods 16, 479–+ (2019).

37. Lonsdale, J. et al. The genotype-tissue expression (GTEx) project. Nat Genet 45, 580–585 (2013).

38. Yang, S.Y. et al. Decontamination of ambient RNA in single-cell RNA-seq with DecontX. Genome Biol 21 (2020).

39. George, J. et al. Comprehensive genomic profiles of small cell lung cancer. Nature 524, 47–53 (2015).

40. Mereu, E. et al. Benchmarking single-cell RNA-sequencing protocols for cell atlas projects. Nat Biotechnol 38, 747–755 (2020).

41. Ritchie, M.E. et al. limma powers differential expression analyses for RNA-sequencing and microarray studies. Nucleic Acids Res 43, e47 (2015).

42. Leek, J.T., Johnson, W.E., Parker, H.S., Jaffe, A.E. & Storey, J.D. The sva package for removing batch effects and other unwanted variation in high-throughput experiments. Bioinformatics 28, 882–883 (2012).

43. Wu, H. et al. Evolution and heterogeneity of non-hereditary colorectal cancer revealed by single-cell exome sequencing. Oncogene 36, 2857–2867 (2017).

44. Wu, H. et al. Single-cell RNA sequencing reveals diverse intratumoral heterogeneities and gene signatures of two types of esophageal cancers. Cancer letters 438, 133–143 (2018).

45. Holik, A.Z. et al. RNA-seq mixology: designing realistic control experiments to compare protocols and analysis methods. Nucleic Acids Res 45, e30 (2017).

